# TOE1 influences canonical Wnt signalling in myeloid leukaemia cells through LEF-1 modulation and regulates the proliferation of haematopoietic cells through PAK2

**DOI:** 10.64898/2025.12.11.691106

**Authors:** H Park, O Sevim, M Wagstaff, A Goff, D Palmer, B Kim, K Heesom, Allison Blair, S Newbury, EL Morgan, BP Towler, TJ Chevassut, RG Morgan

## Abstract

Acute myeloid leukaemia (AML) is an aggressive haematological malignancy characterised by the clonal proliferation of myeloid progenitor cells in the bone marrow and peripheral blood. Dysregulation of the Wnt/β-catenin pathway has been implicated in the establishment and maintenance of leukaemic stem cells in AML, where higher expression of β-catenin promotes clonogenic capacity, drug resistance and inferior survival. The finding that low levels of Wnt signalling are necessary to maintain normal haematopoiesis makes β-catenin an attractive therapeutic target; however, drug design has been hampered by a poor understanding of its molecular interactions in leukaemia cells. To address this, we previously characterised the β-catenin interactome in myeloid cells and identified a plethora of novel interacting proteins. One such interactor was Target of EGR1 (TOE1), a member of the Asp-Glu-Asp-Asp (DEDD) family of deadenylases with previously uncharacterised function in haematopoietic cells. The β-catenin:TOE1 interaction was detected in the nuclear and cytosolic compartments of myeloid cell lines and primary AML samples, and β-catenin depletion was found to promote the cytosolic accumulation of TOE1. Furthermore, TOE1 levels were found to be overexpressed in primary AML blasts versus normal cord-blood derived CD34^+^ haematopoietic stem and progenitor cells (HSPCs), suggesting that TOE1 levels may be dysregulated in leukaemia. TOE1 depletion abrogated Wnt signalling capacity (TCF/LEF activity), potentially via reduced stability/translatability of the Wnt transcription factor lymphoid enhancing factor 1 (LEF-1). TOE1 depletion further suppressed the proliferation and survival of myeloid leukaemia cell lines (HEL and OCI-AML2) and primary human CD34^+^ HSPC; however, this could not be fully explained through LEF-1 alone, since OCI-AML2 do not express LEF-1. Using tandem mass tag (TMT)-labelling coupled to mass spectrometry analysis in TOE1 deficient HEL and OCI-AML2 cells, we identified and validated p21 (RAC1) activated kinase 2 (PAK2) as a downregulated target that could reduce the proliferation (but not survival) of AML cell lines. Interestingly, ectopic expression of PAK2 was able to partially rescue the proliferation defect in TOE1 depleted myeloid cell lines and primary human CD34^+^ HSPC. In summary, these data reveal TOE1 as novel interacting partner for β-catenin in haematological cells capable of modulating Wnt signalling output via LEF-1, and as a novel mediator of growth and survival in human HSPC and AML cells partly through PAK2 regulation.

## Introduction

Acute myeloid leukaemia (AML) is an aggressive clonal disorder of haematopoietic stem/progenitor cells (HSPC) resulting in arrested myeloid development and enhanced self-renewal properties. Despite the emergence of several new targeted therapies,^1^ the prognosis for most patients remains poor and the heterogeneity of AML demands a larger range of targeted molecular therapies with broad applicability. Dysregulated signal transduction is a hallmark of AML biology and frequently the target of novel therapy design as evidenced by novel agents targeting FLT3 and Hedgehog signaling.^2^ Canonical Wnt/β-catenin signalling is important for the maintenance and development of HSPC,^3–7^ and is frequently deregulated in multiple AML subtypes,^8^ where it sustains leukaemia stem cell (LSC) activity,^9–11^ and drug resistance.^12,13^ Efforts to pharmacologically target the central mediator β-catenin through its stability or transcriptional activity has shown promise to date,^8^ but has been hampered by incomplete understanding of β-catenin’s context-specific interactions.

Our previous interrogation of the β-catenin interaction network in haematopoietic cells revealed new interaction partners such as WT1,^14^ and MSI2,^15^ as part of a dense network of RNA-binding proteins (RBP) and subsequently enriched mRNA with β-catenin,^15,16^ suggestive of a novel post-transcriptional role in haematological cells.^17^ One such novel partner identified was **t**arget **o**f **E**GR1 (TOE1), a 510-amino acid member of the Asp-Glu-Asp-Asp (DEDD) family of deadenylases with previously uncharacterised function in haematopoietic cells. The TOE1 gene was only identified and characterised relatively recently, being identified as a suppressor of cell growth mediated by EGR1 via induction of the cell cycle regulator p21.^18^ Subsequent studies have more definitively characterised TOE1 as a Cajal body localised protein with 3’ exonuclease function controlling the maturation/stability of small nuclear RNAs (snRNA),^19–22^ and disruption of this function through biallelic loss-of-function *TOE1* mutations was identified as a fundamental driver of the neurodegenerative syndrome Pontocerebellar Hypoplasia type 7 (PCH7).^23–25^ Additional studies have begun to unravel the complexity of TOE1 function with diverse roles (RNA related or not) reported in telomere maintenance,^26^ p53 transcriptional activity,^27^ and inhibition of HIV transcription and replication in infected T-cells.^28^

The role of TOE1 in human cancer or stem cell biology remains underexplored. Of the limited studies available, one report in gastric cancer indicates a tumour suppressor role for TOE1,^29^ whilst in hepatocellular carcinoma TOE1 may serve oncogenic roles,^30^ particularly through a MYC-STAMBPL1 axis promoting EGFR stability and subsequent Lenvatinib sensitivity.^31^ Indeed, small molecule inhibitors for TOE1 have been screened and predicted to have anti-cancer activity.^32^ The aim of this study was to assess any molecular crosstalk between β-catenin and TOE1 in myeloid cells, and characterise the wider role of TOE1 in a haematopoietic setting for the first time.

## Method

### Primary samples

Bone marrow, peripheral blood or leukapheresis samples from patients diagnosed with AML (clinical information provided in Supplemental Table S1) were collected in accordance with the Declaration of Helsinki and with approvals of University Hospitals Bristol NHS Foundation Trust and London Brent Research Ethics committee. Human cord blood (CB) was obtained following informed consent from healthy mothers at full-term undergoing elective caesarean sections at Royals Sussex County Hospital and Princess Royal Hospital, with approval from University Hospitals Sussex NHS Foundation Trust, the East of England-Essex Research Ethics Committee, HRA and Health and Care Research Wales (18/EE/0403). Mononuclear cells (MNCs) with viability >80% following isolation via density gradient separation using Ficoll-Hypaque (Merck-Millipore, Gillingham, Dorset) were included in the study. The CD34^+^ fraction was enriched as previously described^33^ from cryopreserved cord blood MNC preparations using MiniMACS (Miltenyi Biotec, Woking, Surrey) according to the manufacturer’s instructions.

### Cell culture and drug treatments

The myeloid cell lines K562, HL60, HEL, U937, PLB-985, NOMO1, OCI-AML3, EOL-1, ML-1, THP-1, KU812 (The European Collection of Authenticated Cell Cultures) and OCI-AML2, MV4;11, KG1, KG1a SET-2, NB4 and Mono-Mac-6 (Leibniz Institute DSMZ-German Collection of Microorganisms and Cell Cultures GmbH) were confirmed mycoplasma-free and authenticated via short-tandem repeat (STR) analysis prior to general culture as previously.^34^ β-Catenin was stabilised utilising a GSK3β inhibitor CHIR99021 (Merck-Millipore) overnight as previously described.^16^ Proteasomal turnover and autophagy were inhibited through the use of 1µM MG132 (Merck-Millipore) and 100nM Bafilomycin A1 (Merck-Millipore) respectively, whilst translation was inhibited with 100nM cycloheximide (Merck-Millipore). Purified human CB CD34^+^ HSPCs were isolated and cultured at 5×10^5^/ml in StemSpan SFEMII (StemCell Technologies, Cambridge, Cambridgeshire) as previously described.^16^

### Whole cell lysis

2-5×10^6^ cells were washed in ice cold PBS and resuspended in 100μL 1x lysis buffer (Cell Signaling Technology, Leiden, Netherlands) containing Complete Mini Protease Inhibitor cocktail (Roche, Welwyn Garden City, Hertfordshire) and incubated for 30min with occasional vortexing (Merck-Millipore) to maximize lysis. Insoluble material was removed by centrifugation at 21,000 x *g* for 10min and resulting homogenate stored at -80°C until further use.

### RT-qPCR

RNA was extracted with the Zymo RNA Miniprep kit (Cambridge Bioscience, Cambridge, Cambridgeshire), subjected to DNase treatment with TURBO DNase^TM^ (Thermo Fisher Scientific, Altrincham, Cheshire) and cleaned with the Monarch® RNA Cleanup kit (New England BioLabs, Hitchin, Hertfordshire) according to the manufacturer’s instructions. All primers used in the study are listed in Supplementary Table S2 and were optimised to show efficiency between 90-110% using a SYBR-green kit (Apto-Gen, London, UK). RT-qPCR programmes used as per manufacturer’s instructions (Merck-Millipore). RT-qPCR analysis was performed on QuantStudio^TM^ 3 Real-Time PCR System (Thermo Fisher Scientific) and analysed on Design & Analysis 2 (Thermo Fisher Scientific). Relative gene expression was calculated with the ΔΔCT method with *GAPDH* as the reference gene.

### TMT Labelling and High pH reversed-phase chromatography

Aliquots of 50µg of each sample were digested with trypsin (1.25µg trypsin; 37°C, overnight), labelled with Tandem Mass Tag (TMT) six plex reagents according to the manufacturer’s protocol (Thermo Fisher Scientific) and the labelled samples pooled. The pooled sample was desalted using a SepPak cartridge according to the manufacturer’s instructions (Waters, Milford, Massachusetts, USA). Eluate from the SepPak cartridge was evaporated to dryness and resuspended in buffer A (20 mM ammonium hydroxide, pH 10) prior to fractionation by high pH reversed-phase chromatography using an Ultimate 3000 liquid chromatography system (Thermo Fisher Scientific). In brief, the sample was loaded onto an XBridge BEH C18 Column (130Å, 3.5 µm, 2.1 mm X 150 mm, Waters, UK) in buffer A and peptides eluted with an increasing gradient of buffer B (20 mM Ammonium Hydroxide in acetonitrile, pH 10) from 0-95% over 60 minutes. The resulting fractions (concatenated into 15 in total) were evaporated to dryness and resuspended in 1% formic acid prior to analysis by nano-LC MSMS using an Orbitrap Fusion Lumos mass spectrometer (Thermo Scientific).

### Nano-LC Mass Spectrometry

High pH RP fractions were further fractionated using an Ultimate 3000 nano-LC system in line with an Orbitrap Fusion Lumos mass spectrometer (Thermo Scientific). In brief, peptides in 1% (vol/vol) formic acid were injected onto an Acclaim PepMap C18 nano-trap column (Thermo Scientific). After washing with 0.5% (vol/vol) acetonitrile 0.1% (vol/vol) formic acid peptides were resolved on a 500 mm × 75 μm Acclaim PepMap C18 reverse phase analytical column (Thermo Scientific) over a 150 min organic gradient, using 7 gradient segments (1-6% solvent B over 1min., 6-15% B over 58min., 15-32%B over 58min., 32-40%B over 5min., 40-90%B over 1min., held at 90%B for 6min and then reduced to 1%B over 1min.) with a flow rate of 300 nl min^−1^. Solvent A was 0.1% formic acid and Solvent B was aqueous 80% acetonitrile in 0.1% formic acid. Peptides were ionized by nano-electrospray ionization at 2.0kV using a stainless-steel emitter with an internal diameter of 30 μm (Thermo Scientific) and a capillary temperature of 300°C. All spectra were acquired using an Orbitrap Fusion Lumos mass spectrometer controlled by Xcalibur 3.0 software (Thermo Scientific) and operated in data-dependent acquisition mode using an SPS-MS3 workflow. FTMS1 spectra were collected at a resolution of 120 000, with an automatic gain control (AGC) target of 400 000 and a max injection time of 100ms. Precursors were filtered with an intensity threshold of 5000, according to charge state (to include charge states 2-7) and with monoisotopic peak determination set to Peptide. Previously interrogated precursors were excluded using a dynamic window (60s +/-10ppm). The MS2 precursors were isolated with a quadrupole isolation window of 0.7m/z. ITMS2 spectra were collected with an AGC target of 10 000, max injection time of 70ms and CID collision energy of 35%. For FTMS3 analysis, the Orbitrap was operated at 30 000 resolution with an AGC target of 50 000 and a max injection time of 105ms. Precursors were fragmented by high energy collision dissociation (HCD) at a normalised collision energy of 60% to ensure maximal TMT reporter ion yield. Synchronous Precursor Selection (SPS) was enabled to include up to 10 MS2 fragment ions in the FTMS3 scan. The mass spectrometry proteomics data have been deposited to the ProteomeXchange Consortium via the PRIDE partner^35^ repository with the dataset identifier PXD070891.

### Data Analysis

The raw data files were processed and quantified using Proteome Discoverer software v2.4 (Thermo Scientific) and searched against the UniProt Human database (downloaded January 2025: 83095 entries) using the SEQUEST HT algorithm. Peptide precursor mass tolerance was set at 10ppm, and MS/MS tolerance was set at 0.6Da. Search criteria included oxidation of methionine (+15.995Da), acetylation of the protein N-terminus (+42.011Da), methionine loss from the protein N-terminus (-131.04Da) and methionine loss plus acetylation of the protein N-terminus (-89.03Da) as variable modifications and carbamidomethylation of cysteine (+57.021Da) and the addition of the TMT mass tag (+229.163Da) to peptide N-termini and lysine as fixed modifications. Searches were performed with full tryptic digestion and a maximum of 2 missed cleavages were allowed. The reverse database search option was enabled, and all data was filtered to satisfy false discovery rate (FDR) of 5%.

### Immunoblotting

Immunoblotting was performed as described previously^16^ with antibodies to β-catenin, TOE1, PARN, LEF-1, TCF-1, TCF-4, Lamin A/C, α-tubulin, β-actin and GAPDH details listed in **Supplemental Table S3**). Densitometric analysis of LEF-1 expression was performed utilising ImageJ software version 1.54 (National Institute of Health, Bethesda, Maryland, USA) normalising to the GAPDH density present within each sample.

### Lentivirus preparation

K562 and HEL cells were lentivirally transduced with the β-catenin-activated reporter (BAR) or mutant ‘found unresponsive’ control (fuBAR) systems as described previously^16^. Expression plasmids utilised for lentiviral transgene expression are outlined in **Supplemental Table S4.**

### Flow cytometry

For Wnt reporter assessment, 2×10^5^ BAR/fuBAR containing cells were treated overnight with either DMSO or CHIR99021. TCF/LEF reporter activity (Venus Yellow Fluorescent intensity) was assessed through CytoFLEX Flow cytometer (Beckman Coulter, Amersham, Buckinghamshire) and analysed with FlowJo software version 10.8.2 (Tree Star Inc., Ashland, OR). For immunophenotyping, up to 5×10^4^ CB-derived CD34^+^ HSPCs were resuspending in 100µL staining buffer (1x OBS, 0.5% BSA) containing 10µg/mL CD34-PE, CD45-PerCPCy5.5, CD36-PE and CD13-PerCPCy5.5 (all Biolegend, London, UK) or the equivalent concentration-, manufacturer- and isotype-matched control antibodies. 7-AAD (Thermo Fisher Scientific) was utilised to exclude non-viable cells according to the manufacturer’s instructions.

### Annexin V/PI assay

The Annexin V Apoptosis Detection Kit (BD Pharmingen) was utilised to quantify the following sub-populations: viable/live (Annexin V^neg^, PI^neg^), early apoptotic (Annexin V^pos^, PI^neg^), late apoptotic (Annexin^pos^, PI^pos^) and necrotic/dead cells (Annexin V^neg^, PI^pos^).

### Immunofluorescence (IF)

2×10^6^ cells of interest were washed with PBS and resuspended in 1mL of 2% paraformaldehyde and incubated for 20 minutes at room temperature with agitation. The cells were subsequently resuspended in 1mL quenching buffer (100mM glycine in PBS) and 1mL permeabilisation buffer (0.1% Triton TX100 in PBS) with further washes with staining buffer (1x OBS, 0.5% BSA). The primary antibodies utilised for IF were: anti-TOE1 (Proteintech, Manchester, UK; Bethyl Laboratories, Montgomery, TX), anti-β-catenin (BD Biosciences, Wokingham, Berkshire). The secondary antibodies utilised were: goat anti-mouse Alexa Fluor 488 and goat anti-rabbit Alexa Fluor 647 (Invitrogen, Paisley, Renfrewshire). DAPI was utilised as the nuclear counterstain. Images were captured on the Zeiss LSM 880 confocal microscope utilising a 63x oil immersion objective with the Zen (Black Edition) software (version 2.3).

#### RIP/CLIP

RIP/CLIP assays were performed as previously^36^.

### Nuclear/cytoplasmic fractionation

Nuclear/cytoplasmic fractionation was performed as previously^16^

### Statistics

The Student’s *t-*test was utilised to analyse differences between control and modulated gene expression conditions utilising three biological replicate samples unless otherwise stated. All graphs were prepared utilising GraphPad Prism (software version 8.0.1; GraphPad, Boston, MA). All data are expressed as the mean ± 1 standard deviation (SD) unless otherwise stated.

## Results

### TOE1 is predominantly nuclear localised in myeloid cells and interacts with cytosolic and nuclear β-catenin

Our previous β-catenin interactome data suggested TOE1 interactions across multiple myeloid cell lines including K562, HEL, THP1 and HL60 (**Figure 1A**), but not SW620 colorectal cancer cells,^16^ implying a potential tissue-dependent interaction. TOE1 also has a low presence on the CRAPome database (31/716 = <5%)^37^ reducing its likelihood of representing a background contaminant and rationalising its further investigation. To select models for onward study we initially performed a screen of TOE1 expression across 18 myeloid cell lines and observed ubiquitous expression across all cell lines alongside its functional homolog PARN, and a high degree of co-expression with β-catenin (**Figure 1B**). To confirm interaction with β-catenin we performed reciprocal TOE1 Co-IPs in K562 and HEL cells and observed consistent enrichment of β-catenin under both basal and Wnt signalling stimulated conditions (**Figure 1C**). Since TOE1 has not been studied in a haematopoietic context, we sought to characterise its subcellular localisation. Through nuclear/cytosolic fractionation assays, we observed TOE1 to be a predominantly nuclear-localised protein (apart from HL60) and its localisation was unaffected by β-catenin stabilisation via GSK3β inhibition (**Figure 1D**). Confocal microscopy confirmed a predominant nuclear TOE1 presence with some intense nucleolar-like foci that could be consistent with its previous reported localisation into Cajal bodies (**Figure 1E**).^19^ β-Catenin levels were elevated in the nucleus upon Wnt signalling activation (**Figure 1F**), and compartment specific TOE1 Co-IP confirmed interaction with β-catenin in both the cytoplasm and nucleus of myeloid cells (**Figure 1G**). Finally, given β-catenin and TOE1’s previous associations with mRNA,^21,25,26,38,39^ we ascertained whether interaction was a consequence of mRNA co-occupancy. Despite complete digestion of RNA in Co-IP input samples via RNaseA treatment (**Figure 1H**), the β-catenin:TOE1 interaction remained in both K562 and HEL cells (**Figure 1I**). Furthermore, AlphaFold modelling also predicted against a direct protein interaction between β-catenin and TOE1 (**Supplemental Figure S1A**) suggesting these proteins could be part of a larger multi-protein complex. Taken together, these data confirm that β-catenin forms an RNA-independent indirect interaction with TOE1 in myeloid cells.

**Figure 1.**
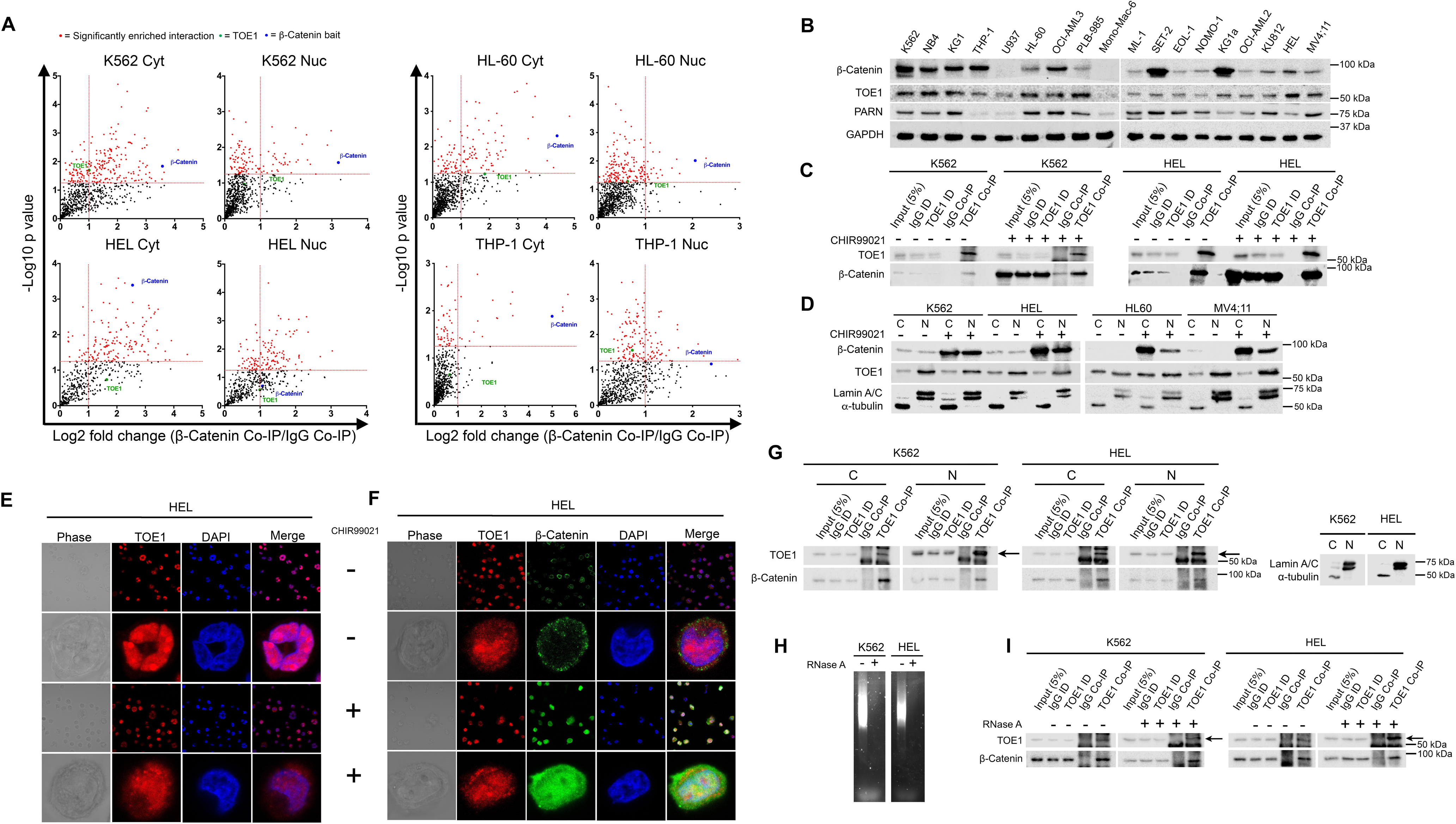
β-Catenin interacts with TOE1 in myeloid leukaemia cells. **(A)** Scatter plots demonstrating β-catenin protein interactions detected in K562 cytosolic/nuclear, HEL cytosolic/nuclear, HL60 cytosolic/nuclear and THP-1 cytosolic/nuclear fractions. The vertical dashed red line indicates the threshold for 2-fold change in protein binding at log2 (=1) in β-catenin Co-IP relative to IgG Co-IP. The horizontal red line represents the threshold for significant interactions at p=0.05 on log_10_ scale (=1.3). Highlighted red dots indicate statistically significant interactions, with TOE1 labelled in green. The remaining black dots represent other proteins detected in the mass spectrometry analysis. Fold change values less than 0 are not shown as these represent isotype Co-IP enriched events. **(B)** Immunoblot demonstrating the endogenous protein expression of β-catenin, TOE1 and PARN across a panel of 18 myeloid cell lines. GAPDH was utilised as the loading control. **(C)** Representative immunoblot showing the level of β-catenin protein present in TOE1 Co-IP derived from K562 and HEL cells under basal (DMSO) versus induced (5µM CHIR99021) Wnt signalling conditions. **(D)** Representative immunoblot showing the protein levels of β-catenin and TOE1 under basal (DMSO) versus induced (5µM CHIR99021) Wnt signalling conditions in K562, HEL, HL60 and MV4;11 cells. Lamin A/C and α-tubulin were utilised to indicate the loading and purity of the nuclear (N) and cytosolic (C) fractions respectively. **(E-F)** Confocal microscopy laser scanning sections demonstrating the sub-cellular localisation of **(E)** TOE1 and **(F)** TOE1 in conjunction with β-catenin under basal (DMSO) versus induced (5µM CHIR99021) Wnt signalling conditions in HEL cells. Phase (grey), TOE1 (red), β-catenin (green), DAPI (blue) and merged TOE1/DAPI images depicted. **(G)** Representative immunoblot demonstrating the level of β-catenin present in TOE1 Co-IP derived from the cytosolic (C) and nuclear (N) fractions of K562 and HEL cells. Lamin A/C and α-tubulin were utilised to indicate the loading and purity of the N and C fractions respectively. Non-specific bands above and below the TOE1 band (57 kDa) were observed in the Co-IP analysis, with the specific TOE1 band represented by an arrow. **(H)** Representative agarose gel electrophoresis images demonstrating RNA in K562/HEL whole cell lysates treated with ± 20mg/mL RNase A overnight prior to TOE1 Co-IP analyses. **(I)** Representative immunoblot showing the level of β-catenin protein present in TOE1 Co-IP derived from K562 and HEL cells ± 20mg/mL RNase A. Non-specific bands above and below the TOE1 band (57 kDa) were observed in the Co-IP analysis, with the specific TOE1 band represented by an arrow.

### TOE1 is overexpressed in AML and associated with poor risk and lower overall survival

Following confirmation of TOE1 interaction with β-catenin in myeloid cells we next explored the likely clinical relevance of TOE1 in AML. Using the adult AML New England Journal of Medicine (NEJM) 2013 dataset from CBioPortal, we found that higher *TOE1* mRNA expression was significantly enriched in poor risk AML patients (**Figure 2A**), and was also linked with reduced overall survival (**Figure 2B**). We next examined the protein expression of TOE1 and β-catenin using a previously interrogated panel of primary AML patient samples.^14^ We observed faint but detectable levels of TOE1 in normal cord blood-derived CD34^+^ HSPC, however TOE1 expression level were higher in 25/28 (89.3%) of AML samples, often with multiple banding (**Figure 2C**). Across the panel, 19/28 (67.9%) samples exhibited co-expression of TOE1 and β-catenin, but there was no correlation in the level of the two proteins (R=0.06, *P*=0.787). Finally, in a primary AML patient sample expressing abundant levels of both β-catenin and TOE1 where ample cellular material existed (patient #20; MLL rearrangement t(9;11) with M5a morphology), we evaluated the β-catenin:TOE1 interaction through TOE1 Co-IP, and observed substantial enrichment of β-catenin (**Figure 2D**). In summary, these data indicate that TOE1 is dysregulated in AML with the potential to impact survival and was thus worthy of further investigation.

**Figure 2.**
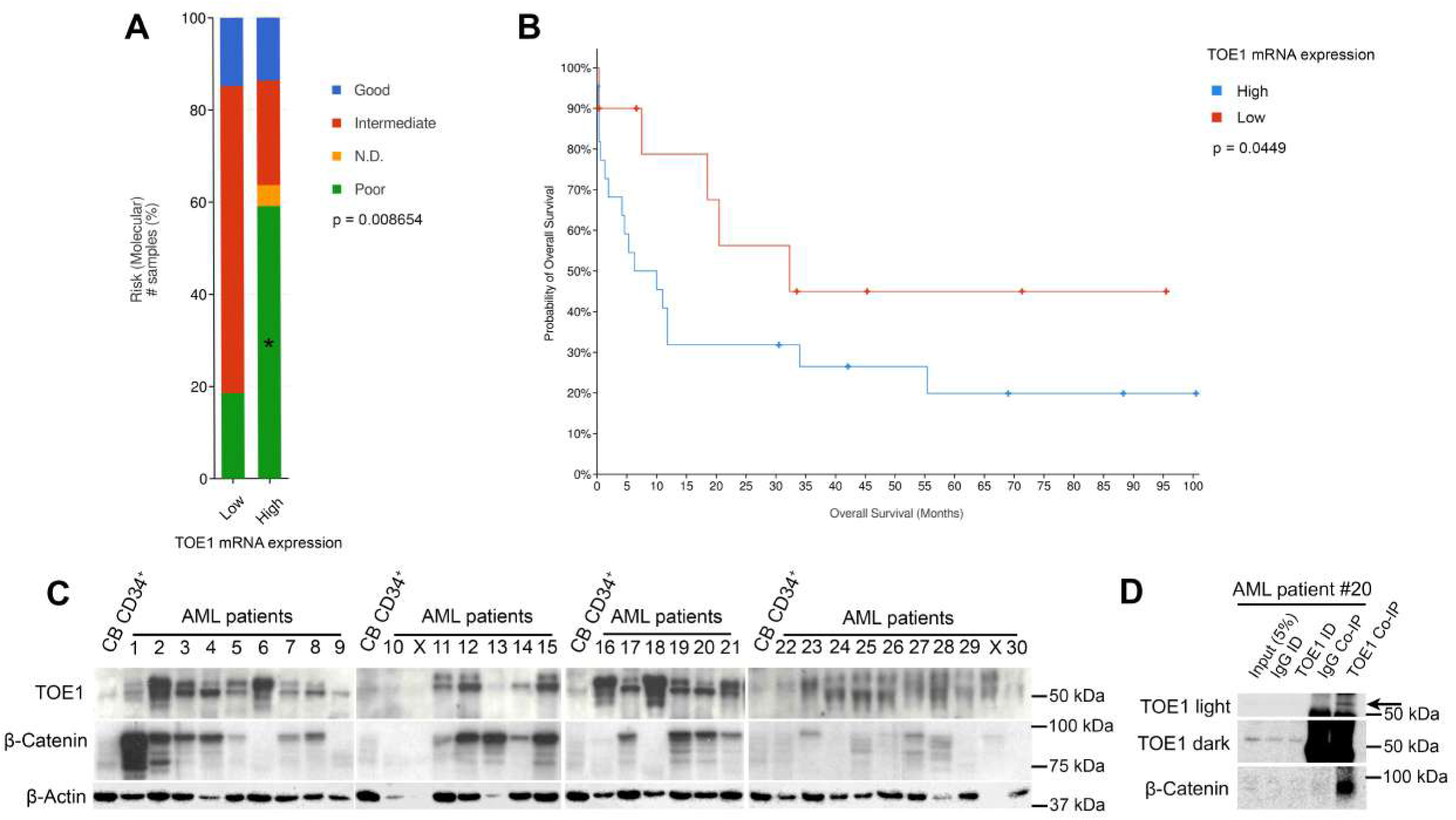
TOE1 is overexpressed in AML and associated with poor risk and lower overall survival. **(A)** Bar chart demonstrating the graphs showing the clinical characteristics of AML patients based on the level of TOE1 mRNA expression. **(B)** Kaplan-Meier curve showing the estimated probability of overall survival in AML patients. TOE1 mRNA levels were derived from the New England Journal of Medicine (NEJM) 2013 The Cancer Genome Atlas (TCGA) whole exome sequencing data set ^60–62^. Low TOE1 mRNA derived via z-score <1; n=27 versus high z score ≥1; n=22. PB, peripheral blood; WBC, white blood cell; N.D, not determined. **(C)** Immunoblot showing the relative levels of β-catenin and TOE1 protein levels in 30 primary AML patient samples alongside a cord blood derived CD34^+^ HSPCs pooled from five independent cord blood samples. X = void samples due to insufficient β-actin levels, which was utilised as a loading control. **(D)** Representative immunoblot showing the level of β-catenin protein present in TOE1 Co-IP derived from AML patient #20 (light and dark exposures shown). Non-specific bands above and below the TOE1 band (57 kDa) were observed in the Co-IP analysis, with the specific TOE1 band represented by an arrow.

### TOE1 regulates Wnt/β-catenin signalling through LEF-1 modulation

Following confirmation of the β-catenin:TOE1 interaction in clinical samples, we next explored the molecular consequence of this relationship through its impact on canonical Wnt signalling activity. To evaluate the role of TOE1 on Wnt signalling output, we generated TOE1 knockdown models in HEL and K562 cells using shRNAs and confirmed reduction of TOE1 protein by immunoblotting which was superior through shRNA#2 (**Figure 3A**). Using the β-catenin-**a**ctivated **r**eporter (BAR) adopted previously,^14,16,34,36^ we first assessed the impact of TOE1 depletion on Wnt signalling output (TCF/LEF activity) and observed a significantly attenuated capacity for Wnt signalling induction in HEL cells in response to the GSK3β inhibitor (and Wnt agonist) CHIR99021 (**Figure 3B**). To understand the underlying cause for this diminished Wnt signalling output we examined expression and nuclear localisation of key Wnt signalling components. We observed no overall change in β-catenin level, or nuclear localisation in response to Wnt agonist treatment (**Figures 3C and D**). We did however observe a significant depletion in the level and nuclear localisation of the Wnt effector LEF-1, which correlated with the efficiency of TOE1 knockdown (**Figures 3C, D and E**).

**Figure 3.**
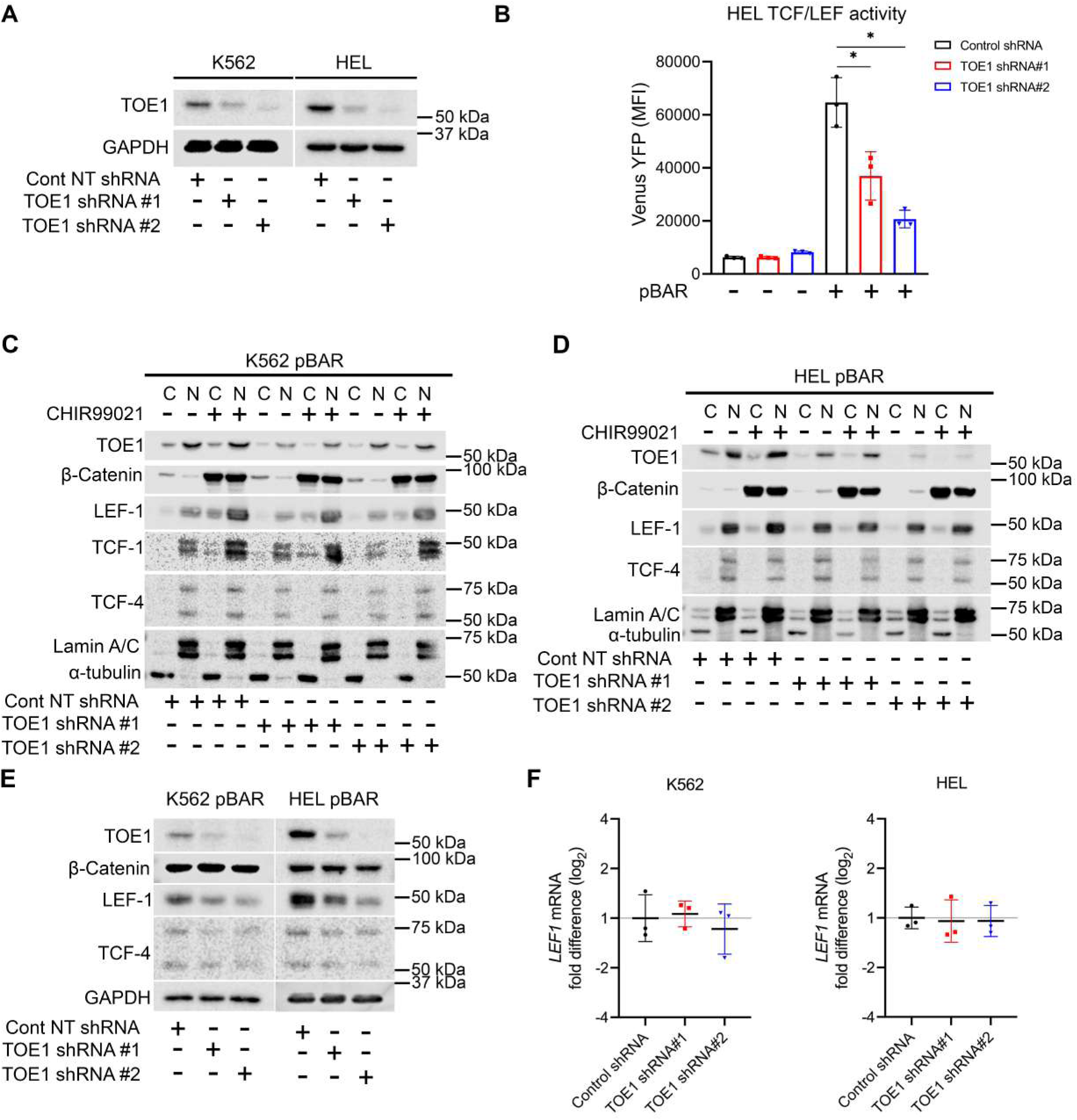
TOE1 regulates Wnt/β-catenin signalling through LEF-1 modulation. **(A)** Immunoblot demonstrating levels of TOE1 expression in and K562 and HEL cells ± TOE1 shRNA. GAPDH was utilised as the loading control. **(B)** Bar graph demonstrating the Venus YFP MFI from the BAR system in HEL cell lines + 5µM CHIR99021 upon TOE1 knockdown (n=3). Error bars indicate mean ± 1SD. Statistical significance is denoted as *p<0.05 (Student’s *t*-test conducted at 72 hours). **(C-E)** Representative immunoblots demonstrating the protein levels of key effector molecules implicated in Wnt signalling ± 5µM DMSO/CHIR99021 in K562 cells and HEL cells ± TOE1 shRNA. HEL cells do not express TCF1. Lamin A/C and α-tubulin were utilised to indicate the loading and purity of the nuclear (N) and cytosol (C) fractions respectively. **(F)** Scatter plot showing the fold change in LEF1 mRNA expression in K562 and HEL cells ± TOE1 shRNA (n=3). GAPDH was utilised as the reference gene (Student’s t- *test*).

Given TOE1’s extensive characterisation as an RNA interacting/modulating protein,^21,23,25,26^ we next sought to ascertain the level of LEF-1 regulation by TOE1 modulation through examination of *LEF1* transcript levels. In both K562 and HEL cells we observed no change in *LEF1* levels in cells harbouring TOE1 shRNA versus non-targeting (NT) shRNA controls (**Figure 3F**). In support of this finding, we also observed no strong association of TOE1 with RNA through both RNA immunoprecipitation (RIP) and crosslinking immunoprecipitation (CLIP) approaches (**Supplemental Figure S1B**), in contrast to a positive control known RNA-binding protein (RBP) LIN28B RIP/CLIP which pulls down abundant RNAs (**Supplemental Figure S1C**). In the absence of *LEF1* mRNA regulation upon TOE1 depletion we instead focussed on protein regulation given TOE1’s recent association with EGFR protein stability.^31^ Our previous data suggests LEF-1 has a very stable half-life,^36^ and have further since confirmed that LEF-1 is not strongly lysosomally regulated (**Supplemental Figure S1D**). However, we have shown that LEF-1 protein stability is modestly controlled by the proteasome in HEL and K562 following 8hrs MG132 exposure in keeping with previous reports (**Supplemental Figure S1E**).^40^ To next ascertain a suitable window for assessing TOE1 impact on LEF-1 peptide stability, we performed a cycloheximide (CHX) chase assay and observed marked LEF-1 degradation following 8hrs of translation inhibition through CHX (**Figure 4A**). Interestingly, TOE1 remained remarkably stable in both K562 and HEL cells even up to maximum 24hrs CHX exposure, suggesting TOE1 is an important protein serving critical housekeeping functions in leukaemia cells. We next examined the capacity of TOE1 to impact LEF-1 protein stability in HEL cells and observed that TOE1 depletion significantly reduced LEF-1 protein stability at 8hrs CHX treatment (**Figure 4B and C**). In summary, these data together demonstrate that TOE1 may regulate Wnt/β-catenin signalling levels through LEF-1 protein stability/translatability.

**Figure 4.**
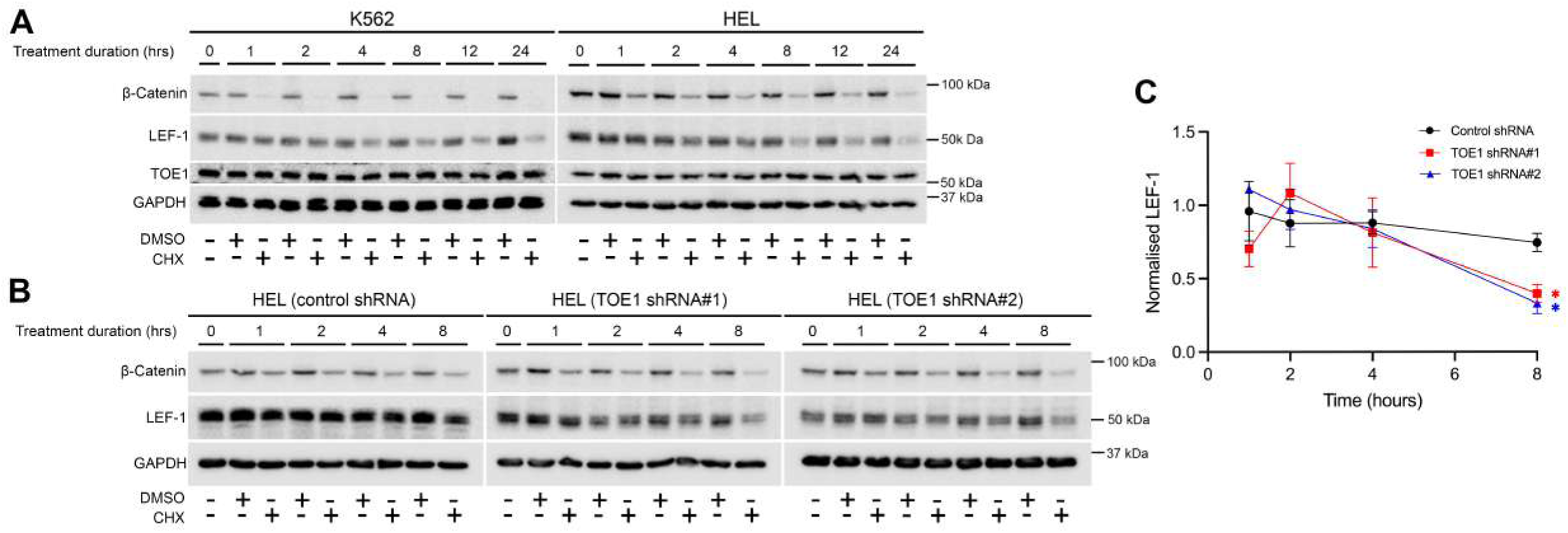
TOE1 regulates LEF-1 protein stability. **(A)** Representative immunoblots showing the protein levels of LEF-1, β-catenin and TOE1 in K562 and HEL cells ± 100nM cycloheximide (CHX). GAPDH was utilised as the loading control. **(B)** Representative immunoblots demonstrating the protein levels of LEF-1, β-catenin and TOE1 in HEL cells ± 100nM cycloheximide (CHX). GAPDH was utilised as the loading control. **(C)** Line graph depicting LEF-1 levels calculated via densitometry analysis, normalised to the relative quantitation values of GAPDH at each time point and DMSO control values (n=3). Errors bars indicate mean ±1 standard error of the mean (SEM). Statistical significance is denoted as *p<0.05 (Student’s t-test).

### TOE1 regulates the growth and survival of haematopoietic cells

Having demonstrated that TOE1 is a β-catenin interacting protein capable of influencing Wnt/signalling output and is dysregulated in AML, we next sought to understand its independent functional roles in AML. Initial exploration into TOE1 function using the DepMap Cancer Dependency Map revealed that many cancer cell lines, including myeloid cells, exhibit TOE1 dependency with largest perturbation effects observed with CRISPR/Cas9 modulation versus RNAi, presumably because of superior target depletion (**Supplemental Figure S2**). Two of the AML cell lines predicted to be sensitive to TOE1 depletion were OCI-AML2 and HEL, therefore we introduced TOE1 shRNA into these lines (**Figure 5A**), and examined their growth over a 72hr culture period. We observed attenuated cell growth rates at 72hrs in both cells lines harbouring TOE1 shRNA versus NT shRNA controls, with the greatest perturbation occurring with the more efficient TOE1 shRNA#2 (**Figures 5B and C**). To examine the cause behind the depleted cell number we examined apoptosis rates using Annexin V staining and observed significantly higher rates of early and late apoptosis across 72 hours with TOE1 shRNA#2 in HEL cells, and significantly higher rates of late apoptosis with both TOE1 shRNAs in OCI-AML2, compared with NT shRNA controls (**Figure 5D and E).** Examination of cell cycle status through DRAQ5 staining revealed no substantial differences between TOE1 versus NT shRNA cells, although larger sub-G0 peaks were observed consistent with a higher rate of apoptosis in TOE1-depleted HEL/OCI-AML2 cells (**Supplemental Figure S3A**).

**Figure 5.**
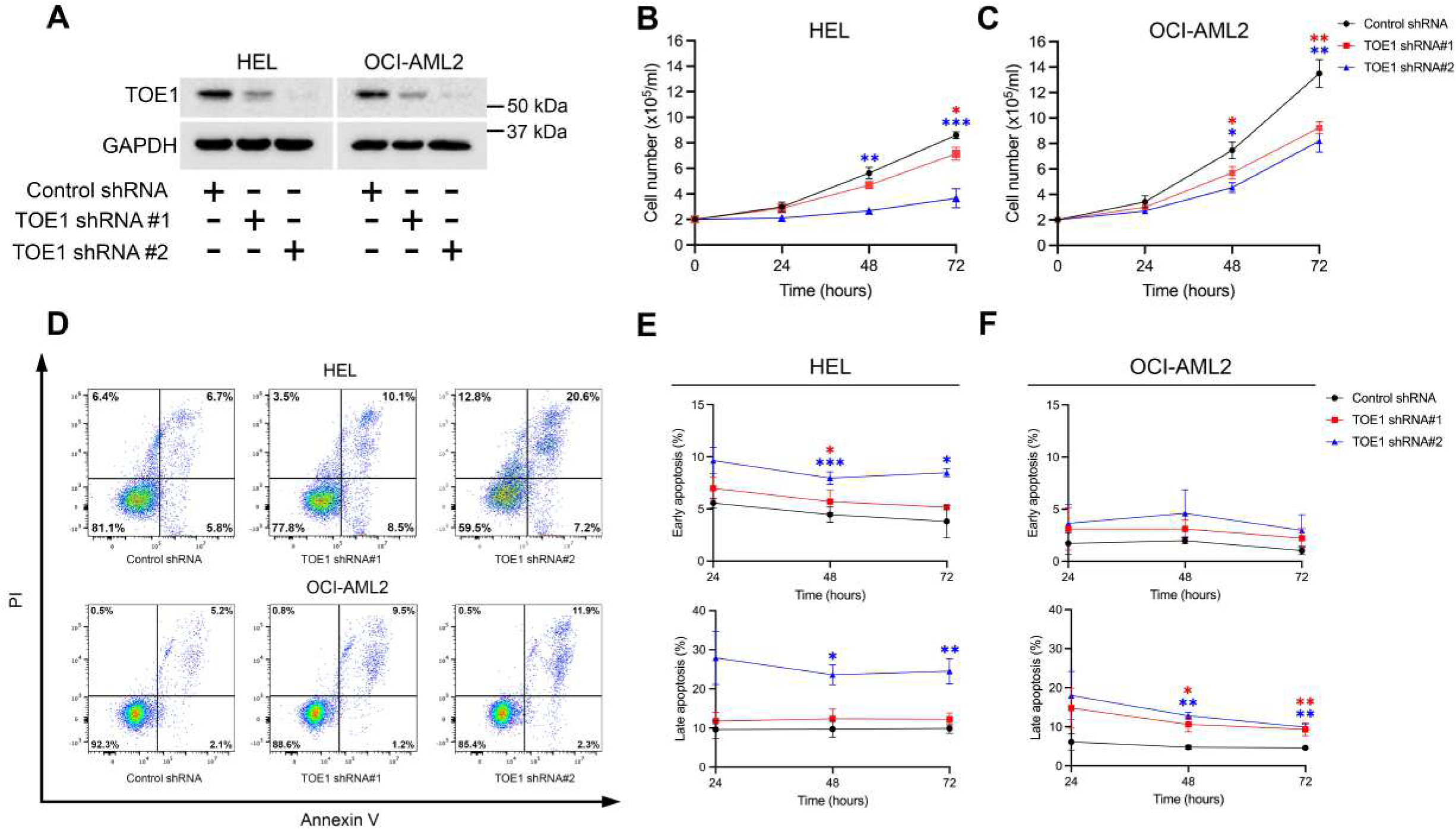
TOE1 impacts the growth and survival of AML cell lines. **(A)** Immunoblot demonstrating levels of TOE1 expression in and HEL and OCI-AML2 cells ± TOE1 shRNA. GAPDH was utilised as the loading control. Line graphs demonstrating the impact of TOE1 depletion on growth of **(B)** HEL and **(C)** OCI-AML2 cells over 72 hours of *in vitro* culture (n=3). **(D)** Representative flow cytometric plots showing cell survival assessed by Annexin V/PI staining in HEL and OCI-AML2 cells ± TOE1 shRNA after 48 hours of *in vitro* culture. The summary data is presented as line graphs over 72 hours (n=3) for **(E)** HEL and **(F)** OCI-AML2 cells. Error bars indicate mean ± 1SD. Statistical significance is denoted as *p<0.05, **p<0.01 (Student’s *t*-test conducted at 72 hours).

Finally, to see if TOE1 regulation of cell growth or survival extended to normal healthy cells we lentivirally transduced normal human CB-derived CD34^+^ HSPC with the optimal TOE1 shRNA#2 containing a GFP selectable marker (**Figure 6A**) and examined both growth rates and broad lineage commitment markers. TOE1 depletion resulted in a gradual depletion in overall GFP^+^ levels from 7 days *in vitro* liquid culture (**Figure 6B**), which resulted in a significant fold-reduction in GFP^+^ levels upon 14-days in steady state differentiation culture (**Figure 6C).** We observed no overt impact of TOE1 on the CD34 positivity of these cultures or broad commitment to monocytic (CD13^+^CD36^+^), granulocytic (CD13^+/-^CD36^-^) or erythroid lineages (CD13^-^CD36^+^) indicating TOE1’s primary influence is over growth/survival, rather than differentiation, of HSPC (**Supplemental Figure S3B**). Taken together, these data indicate that TOE1 positively regulates the growth and survival of both immortalised AML cell lines and normal healthy human HSPC.

**Figure 6.**
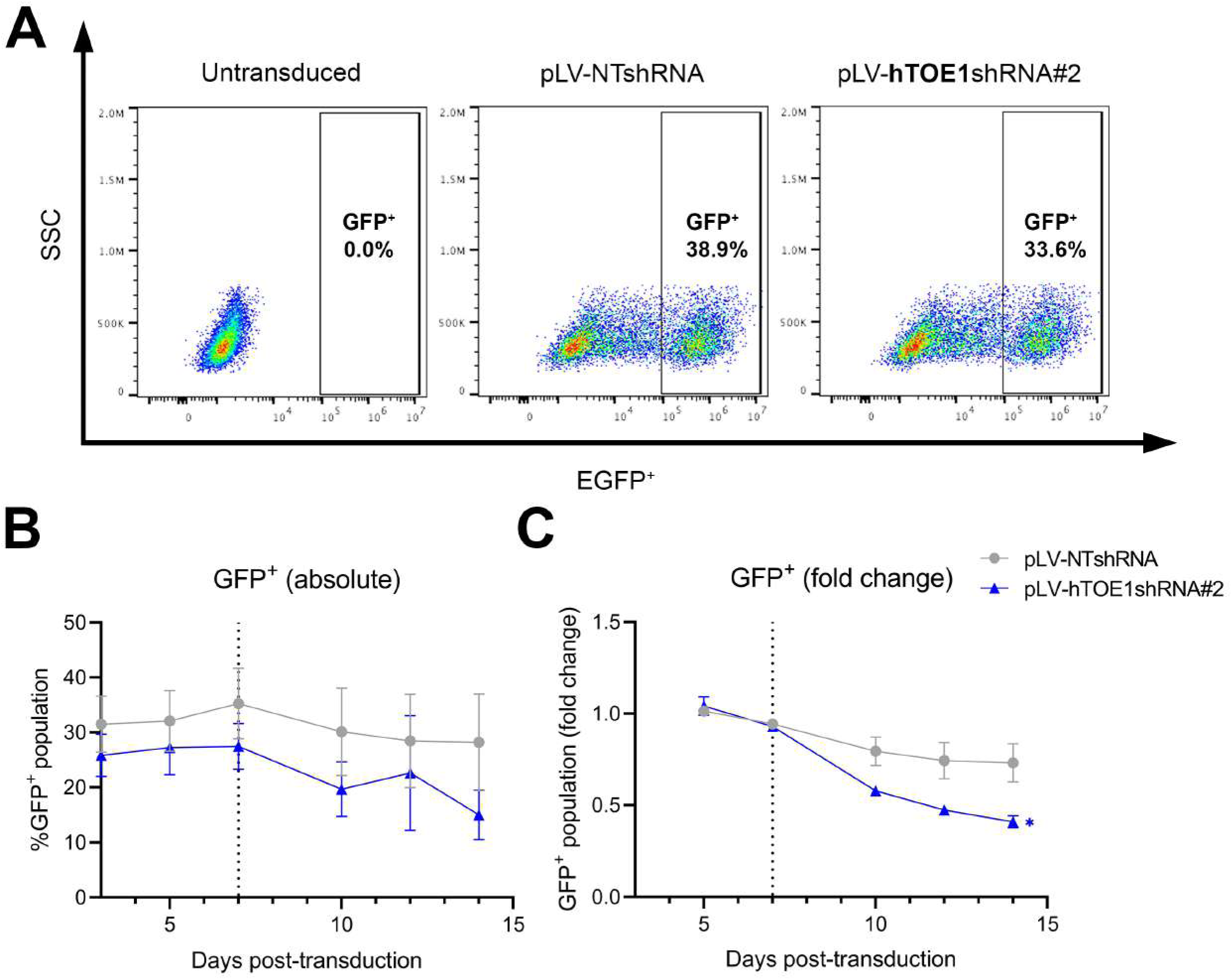
TOE1 knockdown perturbs the proliferation of human CD34^+^ HSPC. **(A)** Representative flow cytometric plots demonstrating the lentiviral efficiency in primary CD34^+^ HSCs 4 days post-transduction. The frequency of GFP^+^ events of human cord blood derived CD34^+^ HSCs transduced with non-targeting shRNA (pLV-NTshRNA) or TOE1 shRNA (pLV-hTOE1shRNA#2) 1 day following isolation, compared to matched untransduced cells. **(B-C)** Line graph showing the absolute and fold change in GFP^+^. Vertical line at day 7 post-transduction represents initiation of steady state differentiation. Error bars indicate mean ± 1SD. Statistical significance is denoted as *p<0.05 (Student’s *t*-test conducted 14 days after lentiviral transduction).

### TOE1 depletion reduces PAK2 protein abundance

Our previous analyses showed that TOE1 could impact LEF-1 expression, and we previously reported that LEF-1 regulates the proliferation of HEL cells,^16^ however we found that OCI-AML2 cells do not express LEF-1 (**Supplemental Figure S4**). Therefore, to uncover a more unifying mechanism for TOE1’s positive regulation of growth/survival in haematopoietic cells we explored the proteome of TOE1-depleted cells using quantitative tandem-mass tag (TMT) labelled mass spectrometry (MS) given we had earlier shown TOE1 has the capacity of influence protein (LEF-1) stability. We generated three experimental replicates for HEL and OCI-AML2 with confirmed TOE1 depletion prior to TMT-labelling and MS analysis (**Figure 7A**). MS data (available via ProteomeXchange with identifier PXD070891) revealed modest but significant alterations to protein abundance between control and TOE1 shRNA cells (**Figure 7B and C**), with 465 proteins downregulated, and 98 upregulated in HEL cells, and 117 downregulated, and 184 upregulated in OCI-AML2 cells (**Figure 7C, Supplemental MS data**). These proteins were associated with a diverse range of biological processes as revealed by Gene Ontology (GO) analysis including many RNA-related functions such as ‘*tRNA modification*’, ‘*translation*’ and ‘*mRNA Pseudouridine synthesis*’, as well as terms pertaining to proliferation including ‘*regulation of mesenchymal stem cell proliferation*’ and ‘*regulation of stem cell proliferation*’ (**Supplemental Figure S5**). Between both cell lines there were a total of 9 commonly downregulated (**Supplemental Table S5**), and 8 commonly upregulated proteins (**Supplemental Table S6**), identified. Given the relatively small number of commonly regulated proteins identified from both cell lines we took a targeted approach to the validation and functional interrogation of putative growth-regulating proteins. Given its known association with growth and survival,^30,41,42^ and the proliferation of murine HSPC,^43^ we selected p21 (RAC1) activated kinase 2 or

**Figure 7.**
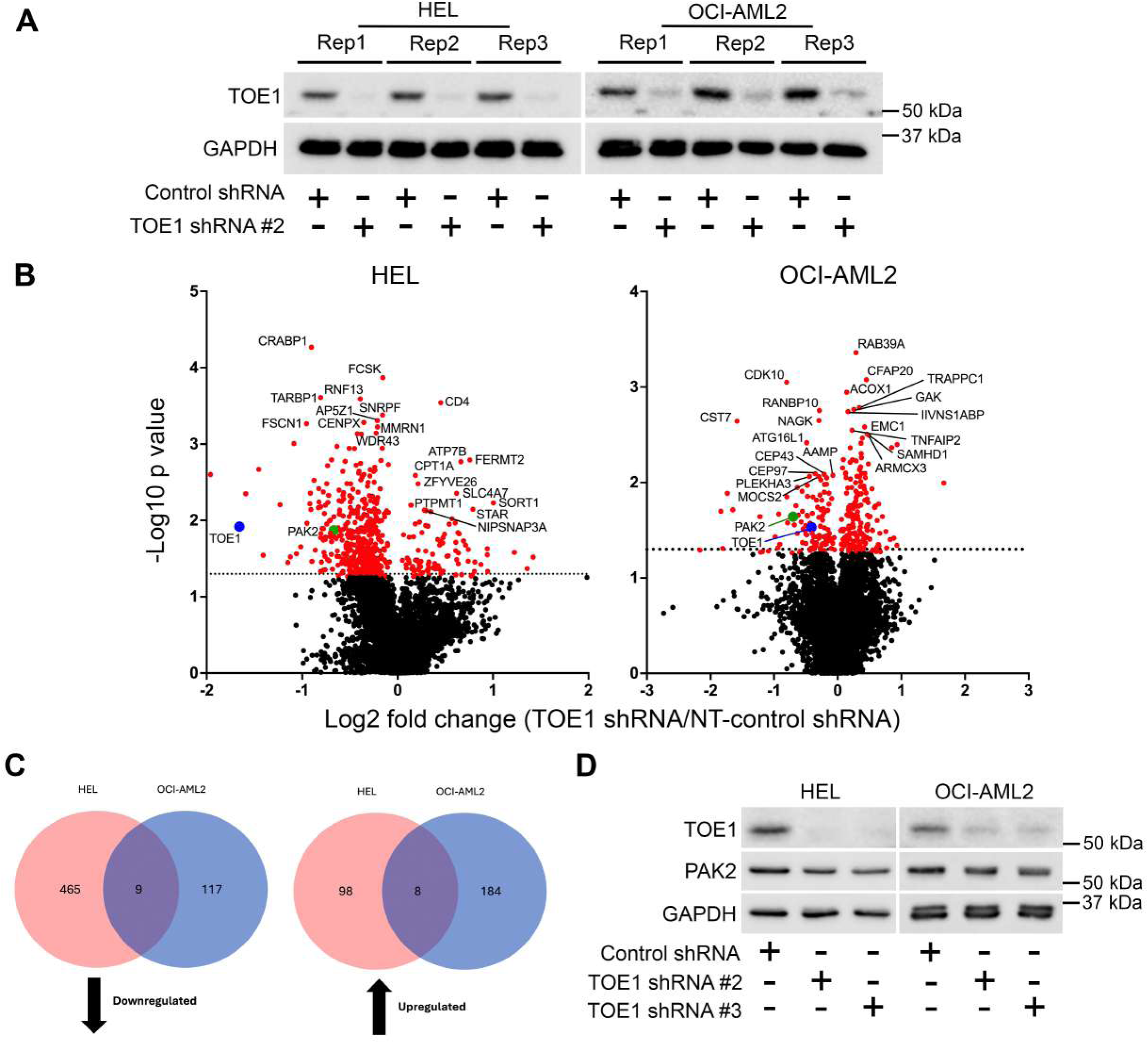
Proteomics analysis of TOE1 depleted AML cell lines identified PAK2 as a commonly regulated protein target. **(A)** Immunoblot demonstrating levels of TOE1 expression in and replicate HEL and OCI-AML2 cells ± TOE1 shRNA prior to mass spectrometry analysis. **(B)** Volcano plots showing differentially expressed proteins in HEL and OCI-AML2 cells with TOE1 shRNA relative to NT-shRNA. Horizonal dashed line indicates the threshold for statistical significance at p*=*0.05 on log_10_ scale (=1.3). Highlighted red dots represent significantly altered peptides (p<0.05). TOE1 is represented as a blue dot whilst PAK2 is highlighted by a green dot. The ten most significantly upregulated and downregulated peptides are annotated for each cell lines. **(C)** Venn diagrams demonstrating the total number of peptides identified to be significantly down- or upregulated in response to TOE1 knockdown, with the common peptides highlighted. **(D)** Representative immunoblot showing the protein levels of PAK2 in HEL and OCI-AML2 cells ± TOE1 shRNA. GAPDH was utilised as the loading control.

Serine/threonine-protein kinase PAK 2 (PAK2) for further investigation. Subsequent validation in HEL and OCI-AML2 via immunoblotting showed a modest but consistent decrease in PAK2 protein expression in response to TOE1 depletion (**Figure 7D**) with stronger regulation also observed in K562 cells (**Supplemental Figure S6**). In summary, these data show TOE1 depletion can modestly impact the cell proteome including regulation of putative growth/survival regulating proteins such as PAK2.

### PAK2 partially mediates the proliferative influence by TOE1 in normal haematopoietic and AML cells

Having shown that TOE1 depletion results in PAK2 reduction, we next interrogated whether this axis could regulate the growth/survival attenuation observed upon TOE1 loss. To address this, we first modulated PAK2 in both HEL and OCI-AML2 using prior-optimised shRNAs (**Figure 8A**) and assessed cell viability and growth. Despite presenting with the most efficient PAK2 depletion, shRNA#1 could not be progressed experimentally since selected cells grew too poorly. As observed in **Figure 8B and C**, PAK2 shRNAs #3 and #4 significantly curtailed cell growth rates in both cell lines but did not impact apoptosis. Since PAK2 appeared to be a positive regulator of proliferation, we next expressed ectopic PAK2 into HEL/OCI-AML2 cells harbouring TOE1 shRNA#2 that were generated previously and examined whether proliferation could be recovered. Given TOE1 shRNA#2 cells were previously selected via puromycin treatment, ectopic PAK2 (prior optimised in **Figure 8A**) was delivered via a GFP selectable construct and following initial lentiviral transduction rates of 60-98% GFP^+^, all double-transduced were enriched to over 98% GFP^+^ via FACS (**Supplemental Figure S7**). Following selection, we next immunoblotted double-transduced cell lines to examine protein expression and confirmed sufficient TOE1 depletion alongside control or ectopic PAK2 levels, whilst also reaffirming PAK2 reduction upon TOE1 depletion (**Figure 8D**). Assessment of proliferation rates showed that single TOE1 shRNA#2 cells exhibited significantly abrogated growth rates as previously observed, which were restored to similar rates as control NT shRNA cells in the presence of ectopic PAK2 (**Figure 8E**). HEL/OCI-AML2 cells harbouring PAK2 overexpression alone did exhibit a small but non-significant increase in proliferation rates. To see if this TOE1:PAK2 regulatory axis could extend to govern the proliferation of healthy cells, we repeated the experiment in normal human CB-derived HSPC. We first observed that TOE1 depletion in puromycin selected primary human HSPC cultures also resulted in PAK2 reduction (**Figure 8F**), indicating this regulatory axis exists in healthy HSPC. Finally, as observed in **Figure 8G**, the presence of ectopic PAK2 was able to overcome the previously observed growth-inhibitory impact of TOE1 depletion in human HSPC over 14 days of *in vitro* culture, and rescue significantly enhanced proliferative expansion. Taken together, these data indicate that TOE1 regulates the proliferation of both normal HSPC, and leukaemia cells, partly through PAK2 modulation.

**Figure 8.**
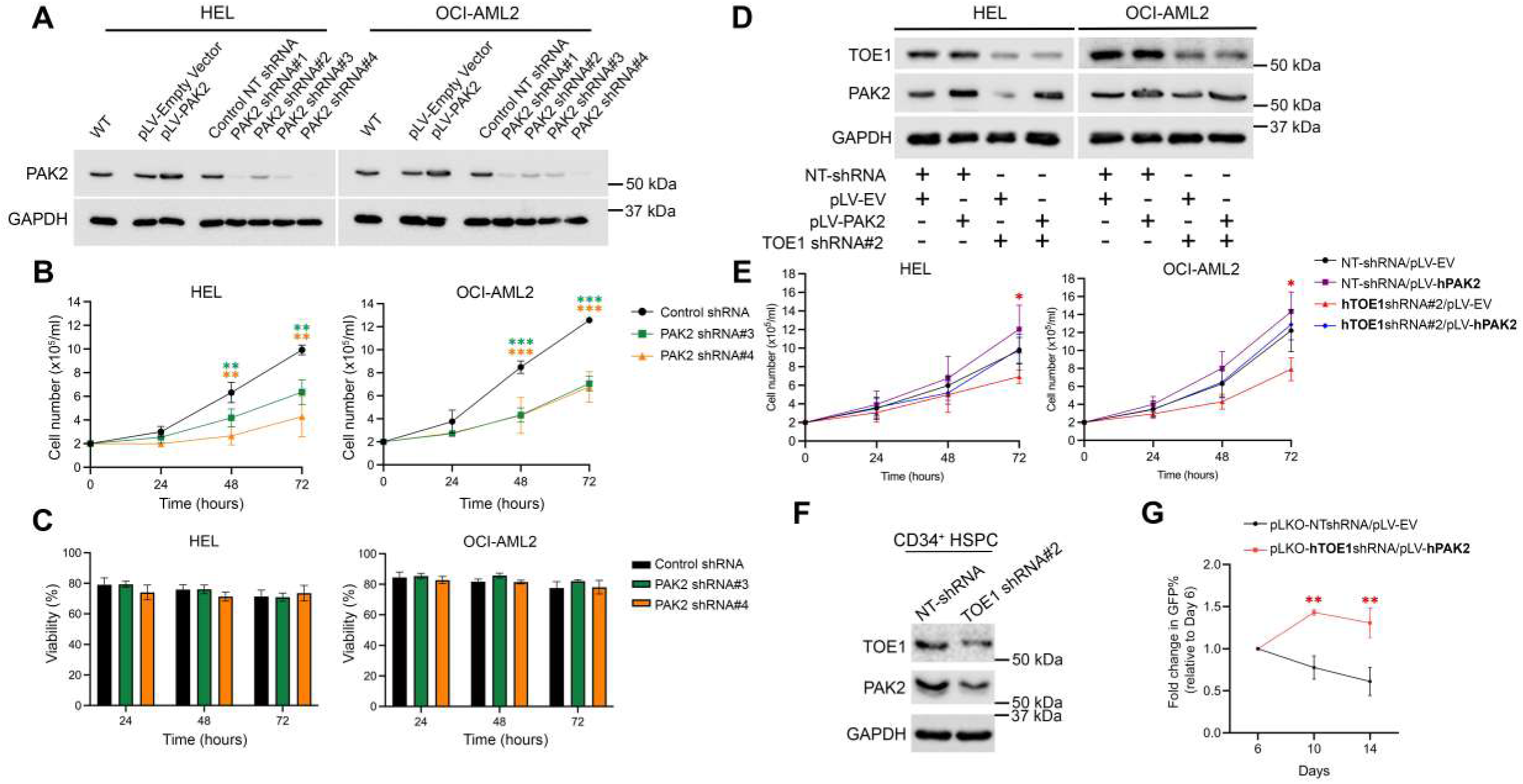
TOE1’s growth promoting influence in leukaemic and normal haematopoietic cells is partly mediated through PAK2. **(A)** Immunoblot showing the levels of PAK2 in myeloid leukaemia cells +/- TOE1 shRNA and +/- pLV-TOE1. GAPDH was utilised as the loading control. **(B-C)** Graphs demonstrating the **(B)** growth and **(C)** viability (calculated via Annexin V/PI staining) of HEL and OCI-AML2 cells ± PAK2 shRNA over 72 hours of *in vitro* culture (n=3). **(D)** Representative immunoblot demonstrating the protein levels of TOE1 and PAK2 following lentiviral transduction with control shRNAs (NT-shRNA or pLV-EV) alongside TOE1 shRNA#2 or pLV-PAK2 and post-FACS purification of EGFP^+^ cells. **(E)** Graphs demonstrating growth over 72 hours of *in vitro* culture under rescue conditions (n=3). **(F)** Representative immunoblot demonstrating the expression of TOE1 and PAK2 in CD34^+^ HSPC ± TOE1 shRNA. **(G)** Line graph showing the fold change in %GFP^+^ cells in CB-derived CD34^+^ HSPC double lentiviral transduced with either NTshRNA/pLV-EV control plasmids or hTOE1shRNA#2/pLV-hPAK2 plasmids and cultured in vitro for 14 days. Error bars indicate mean ± 1SD. Statistical significance is denoted as *p<0.05, **p<0.01, ***p<0.001 (Student’s *t-*test conducted at 72 hours).

## Discussion

This study has uncovered TOE1 as a critical regulator of human leukaemia cell growth/survival for the first time. We found protein levels to be dysregulated across a panel of primary AML samples, and higher *TOE1* (mRNA) expression is associated with inferior patient survival and poor risk disease. To date, TOE1 has been associated with a highly diverse range of cellular functions including telomere maintenance,^26^ p53 regulation,^27^ and viral infection^28^ and our latest data support an emerging role in human cancer. A recent study in hepatocellular carcinoma (HCC) implicated TOE1 in a drug resistance mechanism where it stabilised EGFR expression through a MYC-STAMBPL1 axis.^31^ However, its action could also be context-dependent given its original characterisation as a growth suppressor,^18^ and more recent association as a tumour suppressor in gastric tumours through positive regulation p53 and p21 expression and control of cell cycle progression.^29^ This is also the first report to demonstrate an important role for TOE1 in a human stem cell system, specifically HSPC. This could partly be through its regulation of Wnt/β-catenin signalling (LEF-1) which we also demonstrate in this paper, given that canonical Wnt signalling is known to regulate multiple mammalian stem cell systems.^44^ In mice, TOE1 has been shown to regulate the proliferation and differentiation of neural progenitors. Through TOE1 depletion and transcriptome analysis the authors demonstrated differential expression of several important cell signalling components including those from Notch, TNF and Wnt signalling.^45^ The importance of TOE1 to normal healthy HSPC, and any wider stem systems beyond, could preclude its use as a therapeutically actionable target given the anticipated cytotoxicity. Regardless, there are no clinically approved direct TOE1 inhibitors even though new targeting strategies are emerging. The Cravatt group recently adopted a chemical base-editing strategy to identify essential targetable cysteines for cancer-dependent proteins.^46^ Of the >1750 proteins identified from DepMap Portal, 270 were defined as ‘Strongly Selective’ indicating restricted dependency in specific cancer cell lines. Of a 12-cell line cohort, the growth of MCC142 (Merkel cell carcinoma) and PANC1005 (pancreatic adenocarcinoma) cells were found to be highly TOE1-dependent, extending the number/type of cancers where TOE1 may serve oncogenic function. From here the authors developed two molecules (WX-02–33 or WX-02–13) targeting Cys80 of TOE1, of which WX-02–33 was found to allosterically inhibit the nuclease activity of TOE1 (whilst promoting TOE1 binding to spliceosome complexes) and perturb the growth of MCC142 cell in competition assays. Using WX-02–33 as a reference point, a more recent study by Kehinde *et al*, explored TOE1 Cys80 targeted molecules further with the development of compound 0462 which exhibited more favourable interaction profiles, covalent binding dynamics, free binding energetics, and per-residue energy contributions.^32^ These studies suggest direct TOE1 targeting could be pharmacologically viable should a suitable and safe therapeutic window be identified.

Our original motivation for investigating TOE1’s role in a haematopoietic setting came from its significant enrichment in the β-catenin interactome of multiple myeloid cell lines, which was subsequently validated through reciprocal TOE1 Co-IPs in cell lines and a primary AML patient sample in this study. Despite this robust RNA-independent interaction, we were unable to demonstrate any substantial impact of TOE1 modulation on the level or localisation of β-catenin itself. We intended to assess the impact of β-catenin on TOE1’s RNA editing/interaction capacity; however, we were unable to isolate any detectable RNA through TOE1 RIP or CLIP assessment which was surprising given TOE1’s well established role as a exonuclease/deadenylase for short nuclear non-coding RNAs.^19–23,25,26^ It may be that these TOE1:snRNAs interactions are too transient to be reliably detected through traditional RIP/CLIP approaches, requiring a more sophisticated approach like HyperTRIBE^47^ or a catalytically dead TOE1 variant, to identify TOE1 RNA targets in β-catenin modulated cells, but this was beyond the resources available to the current study. Instead, we opted to focus on the one consistent area of crosstalk we were able to identify between TOE1 and Wnt TCF/LEF activity which was through LEF-1 regulation. We observed no alteration to the overall *LEF1* transcript level and instead found TOE1 altered the stability of LEF-1 as assessed by a cycloheximide chase. Given TOE1’s role in regulating snRNA maturation, we can’t eliminate the possibility that the reduced LEF-1 protein stability upon TOE1 loss isn’t also due to unstable *LEF1* mRNA. Indeed the MYC-STAMBPL1-TOE1 axis which was reported to regulate EGFR expression in HCC, appeared to impact both transcript and protein levels of EGFR which contributed to Lenvatinib resistance.^31^ Part of this mechanism was proposed to be through STAMBPL1’s deubiquitination of TOE1 preventing its lysosomal degradation. However, our experiments in leukaemia cells were unable to demonstrate any strong regulation of TOE1 protein in response to autolysosomal or proteasomal inhibition with Bafilomycin A or MG132, respectively, where it remained remarkably stable. Therefore, the route through which TOE1 regulates the stability of LEF-1 remains unclear currently.

Regardless, LEF-1 alone couldn’t universally explain the reduced growth/survival observed upon TOE1 depletion in AML cells lines/HSPC, since OCI-AML2 do not express LEF-1. To address this, we assessed global protein abundance in TOE1 depleted leukaemia cells for the first time using quantitative mass spectrometry. From this analysis we observed only moderate alterations to global peptide abundance in response to TOE1 depletion. This could possibly be due to incomplete TOE1 ablation, and/or compensation by the functional homolog PARN which we also found to be abundant in myeloid cells (Figure 1A).^21,48^ Throughout our study, we found the degree of TOE1 depletion to be critical in the resulting phenotypes or impact on target expression with better knockdowns demonstrating greater impacts in a dose-dependent fashion. This is reflected in DepMap Portal data where CRISPR/Cas9 mediated TOE1 depletion is more detrimental to cell growth/survival than RNAi, presumably due to superior TOE1 ablation. Regardless, using shRNA we were able to identify significant and consistent alterations to several proteins across both TOE1-depleted AML cell lines, including PAK2. PAK2 was reduced by TOE1 depletion in both AML cell lines and primary *in vitro* cultured HSPCs. PAKs are a family of important evolutionary conserved serine/threonine kinases implicated in numerous critical cellular processes including cytoskeletal arrangement, motility, apoptosis, proliferation and cell division.^49–51^ PAK2 was prioritised for further study given its previous association with several processes important to murine HSPC biology including homing/migration,^52^ and more importantly for this study, proliferation.^43^ HSCs from conditional PAK2 knockout mice exhibited growth and survival defects in multi-cytokine supplemented *in vitro* culture, albeit with no difference in self-renewal capacity, a similar phenotype we observed with TOE1 shRNA in human HSPCs. Our data indicate PAK2 is also important for human HSPC proliferation and indeed AML cell growth more generally, although ectopic PAK2 expression didn’t appear to impact cell viability/survival, implying other mechanisms beyond PAK2 contribute to this phenotype. Elsewhere PAK2 has been implicated in haematological malignancies,^53^ and further human cancers beyond.^54^ Whereas, PAK1 and 4 are the predominant isoforms deregulated in solid tumours,^51^ PAK1 and 2 are key drivers of BCR-ABL+ chronic myeloid leukaemia cells. PAK2, rather than PAK1 was a key regulator of CML cell growth, but only a limited impact on survival was observed until PAK1 and 2 were ablated in combination.^55^ No PAK2 specific inhibitors have been developed yet, however pan-PAK inhibitors such as PF-3758309, FRAX-486 and IPA-3 are available and have shown promise in lymphoid malignancies^56–58^ and AML.^59^ However, pre-existing reports alongside our latest study suggest a very carefully defined therapeutic window is necessary given the likely HSPC toxicity.

In summary, this study has shown for the first time that the β-catenin interacting protein TOE1 impacts Wnt signalling in leukaemia cells through LEF-1 modulation, and its levels are dysregulated in leukaemia where it regulates the proliferation of human AML cells and HSPC through PAK2.

## Author contributions

HP and OS performed experiments, analysed data and co-wrote the manuscript. MW executed experiments and provided laboratory support, whilst AG and DP optimised CLIP assays. BK assisted with cord blood collection/processing, KH performed TMT-LC/MS analysis and AB provided primary AML samples. SN, ELM and BT provided experimental guidance, equipment and reagents. TJC and RGM performed experiments, analysed data, co-wrote the manuscript, secured funding and directed the study.

## Acknowledgements

This work was funded by the Kay Kendall Leukaemia Fund (RM: KKL1051/KKL1446), Leukaemia & Myeloma Research UK (RM: 4-5/06.21R), Children’s Cancer & Leukaemia Group (CCLGA 2023 16 Morgan), British Society for Haematology (HP: 44299), the Sussex Cancer Fund (HP) and the Republic of Türkiye Ministry of National Education (OS). Thanks to Dr Paraskevi Diamanti (University of Bristol) for AML patient sample collection and Lizzy Hoole (Institute for Child Life & Health, UHBristol & Weston NHS Foundation Trust) for supplying clinical data. Thanks to the midwives and clinical research nurses at University Hospitals Sussex (UHS) NHS trust including Raquel Akieme, Valentina Toska, Lorraine Shah-Goodwin, Denise Skinner, Carla Clegg, Edina Lalu and Elohor Uwadiogbu, for the collection of human umbilical cord blood. We thank all the technicians in the School of Life Sciences that maintain our laboratories and facilities. We are indebted to the patients and their families who gave consent for their samples to be used for our research.

